# Nicotinamide Riboside Attenuates inflammation in a mouse model of psoriasis

**DOI:** 10.64898/2026.11.13.766424

**Authors:** Aurea Oliva, Ignacio Cayuela, Jorge S. Yebra, Yara Cuesta, Michael N. Sack, María N. Navarro, Javier Traba

## Abstract

Psoriasis is a chronic inflammatory skin disease driven by dysregulated immune responses, in which IL-17-producing T cells play a central role. Nicotinamide riboside (NR), a precursor of NAD^+^, has recently emerged as a potential immunomodulatory intervention. Previous studies have shown that NR preferentially suppresses Th17 responsiveness in CD4^+^ T cells from individuals with psoriasis, but whether NR can attenuate psoriasiform inflammation *in vivo* remains poorly defined. Here, we investigated the effects of NR supplementation in a mouse model of imiquimod (IMQ)-induced psoriasiform skin inflammation. NR treatment markedly attenuated IMQ-induced splenomegaly and reduced epidermal thickening, indicating an improvement of both systemic and cutaneous inflammatory manifestations. Flow cytometric analysis of skin infiltrates revealed a reduction in inflammatory myeloid populations, including neutrophils and inflammatory monocyte–macrophages, together with a pronounced decrease in dermal TCR γδ^+^ and CD4^+^ T cells. Consistent with these findings, CD4^+^ T cells isolated from IMQ-treated mice displayed increased IL-17A production following *ex vivo* TCR stimulation, which was significantly reduced by NR supplementation in a dose-dependent manner. IFNγ and IL-4 responses were not significantly altered by NR in this setting. Furthermore, NR suppressed IL-17A production in Th17-polarized CD4^+^ T cells, while leaving Th1 and Th2 responses largely unaffected. Together, these findings show that NR treatment attenuates IMQ-induced psoriasiform skin inflammation *in vivo* and preferentially modulates IL-17A-associated CD4^+^ T-cell responses. The accompanying reduction in dermal γδ T-cell numbers suggests that NR may also influence innate-like T-cell populations involved in cutaneous inflammation, providing experimental evidence for NAD^+^ boosting as a potential immunomodulatory strategy in psoriasis.

## INTRODUCTION

Psoriasis is a chronic immune-mediated inflammatory skin disorder characterized by dysregulated crosstalk between innate and adaptive immune compartments, leading to keratinocyte hyperproliferation and persistent cutaneous inflammation. Although CD4^+^ Th17 cells and the IL-23/IL-17 axis are central to disease pathogenesis, other immune populations also contribute to the initiation and amplification of skin inflammation. In particular, innate-like T-cell subsets contribute significantly to early inflammatory responses in the skin (Mills et al., 2023).

Among these populations, γδ T cells have emerged as key drivers of cutaneous inflammation in murine models of psoriasis-like inflammation. Dermal γδ T cells can rapidly respond to inflammatory cues and contribute to the early production of IL-17 and related cytokines, promoting keratinocyte activation and epidermal hyperplasia. γδ17 cells have also been identified in human psoriatic skin lesions. Thus, understanding how metabolic signals influence both adaptive and innate-like IL-17-producing populations may provide further insight into the regulation of psoriatic inflammation.

Recent advances in immunometabolism have highlighted the importance of cellular metabolic programs in regulating immune cell activation and effector function. Nicotinamide adenine dinucleotide (NAD^+^) has emerged as a central metabolic node controlling redox homeostasis, mitochondrial function, and transcriptional regulation (Canto et al., 2015). Nicotinamide riboside (NR), a vitamin B3 derivative and NAD^+^ precursor, can increase intracellular NAD^+^ availability and has therefore attracted interest as a potential strategy to modulate inflammatory responses.

In this context, we previously showed that NR supplementation suppresses inflammatory responses in human CD4^+^ T cells, with particularly pronounced effects on Th17 responses. Using primary CD4^+^ T cells from healthy individuals and patients with psoriasis, we found that NR reduced the production of IL-17 and IFN-γ, with particularly pronounced effects on Th17-associated responses. Mechanistically, these effects were associated with activation of NRF2-dependent antioxidant responses, increased SQSTM1/p62 expression and reduced reactive oxygen species (ROS). Integrated transcriptomic and metabolomic analyses further implicated changes in arginine and fumarate metabolism in the regulation of T-cell inflammatory activity (Han et al., 2023).

More recently, a randomized, placebo-controlled study of oral NR supplementation in individuals with mild-to-moderate psoriasis showed that four weeks of supplementation increased whole-blood NAD^+^ levels and attenuated Th1 and Th17 responses in circulating CD4^+^ T cells. Mechanistically, the study identified the SLIT2–ROBO1 signaling pathway as a mediator of these effects. NR increased SLIT2 production by dermal fibroblasts, while SLIT2–ROBO1 signaling inhibited Rho GTPase activity and Th17 polarization. These findings revealed a link between NAD^+^ metabolism, dermal fibroblast signaling and T-cell responses in psoriasis (Han et al., 2026). Importantly, the study focused on systemic immune responses and cellular mechanisms rather than changes in cutaneous inflammation following NR supplementation.

Despite these advances, it remains unclear whether NR-mediated NAD^+^ augmentation can modulate inflammatory responses within the skin *in vivo*, particularly the accumulation of innate immune populations and innate-like IL-17-producing lymphocytes. Whether NR influences cutaneous γδ T-cell populations, which may contribute to early inflammatory amplification in psoriasis-like disease, has not been established.

In this study, we investigate the effects of NR supplementation in a murine model of imiquimod (IMQ)-induced psoriasis-like skin inflammation. We show that NR administration attenuates IMQ-induced psoriasiform skin inflammation, reducing epidermal thickening and inflammatory cell infiltration. Notably, NR administration was associated with reduced numbers of dermal γδ T cells, together with changes in other myeloid and lymphoid populations. These findings complement previous studies on NAD^+^ augmentation in psoriasis by providing evidence that NR can attenuate cutaneous inflammation in an experimental model of psoriasiform skin inflammation.

## MATERIALS AND METHODS

### Mouse models

All mice used in this study were on a C57BL/6J genetic background. Mice were maintained under specific pathogen-free (SPF) conditions in the animal facility of the Centro de Biología Molecular Severo Ochoa. Mice breeding and experimental procedures were conducted in accordance with national and institutional guidelines for animal care, including EU Directive 2010/63/EU on the protection of animals used for scientific purposes. All experimental procedures were approved by the Comunidad de Madrid (approval no. PROEX 220.8/23).

### CD4^**+**^ T cell isolation and stimulation

Mouse CD4^+^ T cells were negatively selected from spleens using the CD4^+^ T Cell Isolation Kit (Miltenyi Biotec), according to the manufacturer’s instructions. Purified cells were cultured in RPMI 1640 supplemented with 25 mM HEPES, 10% heat-inactivated fetal bovine serum (FBS), and penicillin–streptomycin.

For activation assays, naïve CD4^+^ T cells (3 × 105 cells/well) were seeded in 96-well plates and stimulated with plate-bound anti-CD3 (5 μg/mL; BioLegend) and anti-CD28 (10 μg/mL; BioLegend) for 3 days in the presence or absence of NR.

For polarization experiments, naïve CD4^+^ T cells (2 × 105 cells/well) were seeded in 96-well plates and differentiated into Th1, Th2, or Th17 cells under lineage-specific polarizing conditions. Th1 differentiation was induced with IL-12 (20 ng/mL) and anti-IL-4 antibody (10 μg/mL). Th2 differentiation was induced with IL-4 (10 ng/mL) and anti-IFNγ antibody (10 μg/mL). Th17 differentiation was induced with IL-6 (20 ng/mL), TGF-β1 (2 ng/mL), IL-1β (10 ng/mL), IL-23 (10 ng/mL), anti-IL-4 antibody (10 μg/mL), and anti-IFNγ antibody (10 μg/mL). Cells were stimulated with plate-bound anti-CD3 and anti-CD28 for 3 days under the respective polarizing conditions, in the presence or absence of NR. All recombinant cytokines and neutralizing antibodies were purchased from PeproTech or eBioscience.

Culture supernatants were collected, centrifuged to remove cellular debris, and stored at −80°C until analysis. IFNγ, IL-4, and IL-17A levels were quantified by ELISA (R&D Systems), according to the manufacturer’s instructions. Cytokine levels were normalized to cell number using the CyQUANT Cell Proliferation Assay (Invitrogen).

### Imiquimod treatment

C57BL/6 mice were treated topically with 5% imiquimod (IMQ) cream (Aldara; Viatris Pharmaceuticals) applied to both ears daily for 6 consecutive days (10 mg per day) to induce psoriasis-like skin inflammation.

To assess the therapeutic effect of nicotinamide riboside (NR) *in vivo*, mice received daily intraperitoneal injections of NR (1000 mg/kg) or vehicle control (PBS), starting 2 days before the first IMQ application and continuing throughout the treatment period until sacrifice. At the experimental endpoint, spleens and ears were collected.

For flow cytometric analysis of skin-infiltrating immune cells, right ears were bisected and finely minced, then enzymatically digested for 45 minutes at 37°C in RPMI 1640 containing Liberase (83 µg/mL; Roche), DNase I (100 µg/mL; Roche), and collagenase IV (0.5 mg/mL; Sigma). The remaining tissue fragments were further mechanically dissociated using 7-mm stainless steel beads (Qiagen) in a TissueLyser LT (20 oscillations per 5 minutes; Qiagen) to maximize cell recovery.

For histological assessment, left ears were immediately fixed in 4% paraformaldehyde, processed by the Histology Facility at the Centro Nacional de Biotecnología (CNB-CSIC), embedded in paraffin, and sectioned at 4–5 µm at 200 µm intervals. Sections were stained with hematoxylin and eosin (H&E). Epidermal thickness was quantified in a blinded manner using ImageJ software (https://imagej.net/ij). Measurements were performed on 8 sections per mouse, with 20 randomly selected fields analyzed per section.

### Flow cytometry

Cells were washed twice in PBS and incubated with Ghost Dye Red 780 (Cytek Biosciences) with Fc block (2.4G2, reference no. 553142; BD Biosciences) for 30 minutes on ice before antibody surface staining for dead cell exclusion. Samples were acquired on a FACSCanto II flow cytometer with DIVA software and on a Aurora 4L spectral analyzer with Spectral Flow software (Cytek Biosciences), then analyzed by FlowJo software (Becton Dickinson). Cells were gated according to their forward-scatter and side-scatter profiles, and dead cells were excluded based on their staining with the viability dye. The following antibodies were used (antibody/clone ID): Ly6C/AL-21-PerCP-Cy5.5, IL-17A/TC11-18H10.1-PE, IFN-γ/XMG1.2-APC (BD Biosciences); CD4/GK1.5-APC/Fire, CD11b/M1/70-PE-Dazzle 594, CD11b/M1/70-PerCP-Cy 5.5, CD27/LG.3A10-PE-Cy7, CD44/IM7-BV570, CD45/30-F11-BV421, CD64/X54-5/7.1-BV711, Ly6C/HK1.4-BV570, TCR γδ/GL3-PE, TCR γδ/GL3-APC, CD80/16-10A1-APC (from BioLegend); RORγt/AFKJS-9-APC (from Invitrogen); and CD3/145-2C11-PE, CD3/145-2C11-PerCP-Cy 5.5, CD44/IM7-PE-Cy7, CD44/IM7-V450, CD45.2/104-PE, and Ly6G/1A8-PE-Cy7 (from Tonbo Biosciences).

### Statistical analysis

Statistical analyses were performed using GraphPad Prism (version 11). Data are presented as mean ± SEM unless otherwise indicated. Comparison between two groups were performed using paired or unpaired two-tailed t-tests as appropriate. One-way ANOVA was used when more than two groups were compared. Two-way ANOVA was applied when two independent variables were analyzed. A p-value of < 0.05 was considered statistically significant.

## RESULTS

### NR supplementation dampens the ex vivo responsiveness of CD4^**+**^ T helper cells

To evaluate the immunomodulatory effects of NAD^+^ boosting in mice, psoriasis-like skin inflammation was induced in mice by daily topical application of IMQ (**Fig 1.A)**. At the experimental endpoint, spleens were collected, and CD4^+^ T cells were isolated to assess T helper (Th) cell activation.

**Figure 1.**
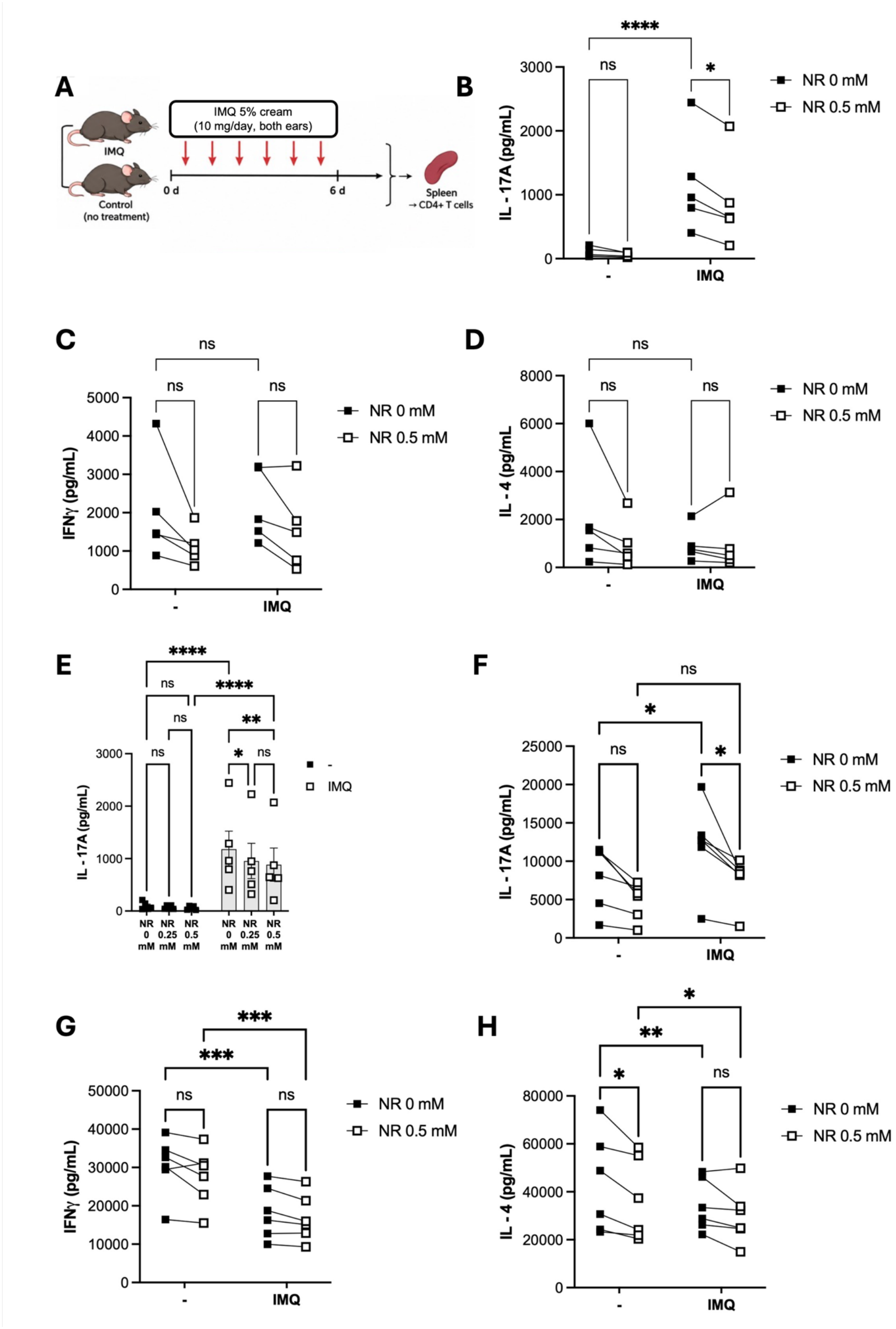
NR supplementation selectively dampens pro-inflammatory CD4+ T-cell responses. (**A**) Experimental design: psoriasis-like skin inflammation was induced by daily topical application of imiquimod (IMQ). At the experimental endpoint, splenic CD4^+^ T cells were isolated and stimulated *ex vivo* through T-cell receptor (TCR) activation. (**B**–**D**) Cytokine production by stimulated CD4^+^ T cells isolated from control or IMQ mice. NR supplementation significantly reduced IL-17A production, whereas IFNγ showed a marked but non-significant reduction. IL-4 production was not altered. Data are shown as mean ± SEM, n=5. (**E**–**G**) Dose-dependent effect of NR on cytokine production by activated CD4^+^ T cells. Increasing concentrations of NR preferentially inhibited IL-17A production compared with IFNγ, while IL-4 secretion remained unchanged. Data are shown as mean ± SEM, n=5. (**H**–**J**) Cytokine production by lineage-polarized Th1, Th2, and Th17 cells cultured in the presence or absence of NR. NR significantly reduced IL-17A secretion by Th17 cells, whereas IFNγ production by Th1 and IL-4 production by Th2 cells was not significantly affected. Data are shown as mean ± SEM, n=6. Statistical significance was determined using two-way ANOVA followed by Tukey’s multiple comparisons test. *p < 0.05, **p < 0.01, ***p < 0.001, ****p < 0.0001.

CD4^+^ T cells isolated from control or IMQ-treated mice were stimulated *ex vivo* through T-cell receptor (TCR) activation. Given that IMQ-induced psoriasis is primarily driven by Th1- and Th17-mediated immune responses, IFNγ and IL-17A production were quantified to assess the effects of NR treatment. As expected, CD4^+^ T cells from IMQ-treated mice produced significantly higher levels of IL-17A than cells from control mice, confirming the induction of a Th17-associated response by IMQ (**Fig 1.B**). IFNγ production showed a slight increase in IMQ-treated mice, although this difference did not reach statistical significance (**Fig 1.C**), whereas IL-4 production showed a slight, non-significant decrease (**Fig 1.D**).

Importantly, NR supplementation significantly reduced IL-17A production by CD4^+^ T cells from IMQ-treated mice (**Fig 1.B**). This reduction was dose-dependent (**Fig. 1E**) and more pronounced than the effect on IFNγ, which showed a marked but non-significant reduction (**Fig 1.C**). In contrast, IL-4 production remained unchanged by NR supplementation (**Fig 1.D**), indicating that Th2 responses were not affected by NR treatment. The effect of NR was dose dependent (**Fig 1.E**). In agreement with the human data (Han et al., 2023), these findings identify the suppression of IL-17A production as the most prominent effect of NR on CD4^+^ T-cell responses, supporting a preferential inhibition of Th17-associated inflammation.

To further investigate this effect, we assessed NR responsiveness in specific lineage-polarized T cells (Th1, Th2 and Th17), where a similar pattern was observed. NR significantly reduced IL-17A production in Th17-polarized cells (**Fig 1.F**). In contrast, IFNγ and IL-4 secretion by Th1- and Th2-polarized cells, respectively, were not significantly altered (**Fig 1.G-H**). These results further support a preferential suppressive effect of NR on Th17-associated IL-17A production, with no significant changes in Th1- or Th2-associated cytokine responses under these polarization conditions.

### NR treatment attenuates imiquimod-induced splenomegaly

To further investigate the effects of NR treatment *in vivo*, we assessed spleen size as an indicator of systemic inflammation in the IMQ-induced psoriasis-like mouse model. The experimental design is summarized in **Fig. 2A**.

**Figure 2.**
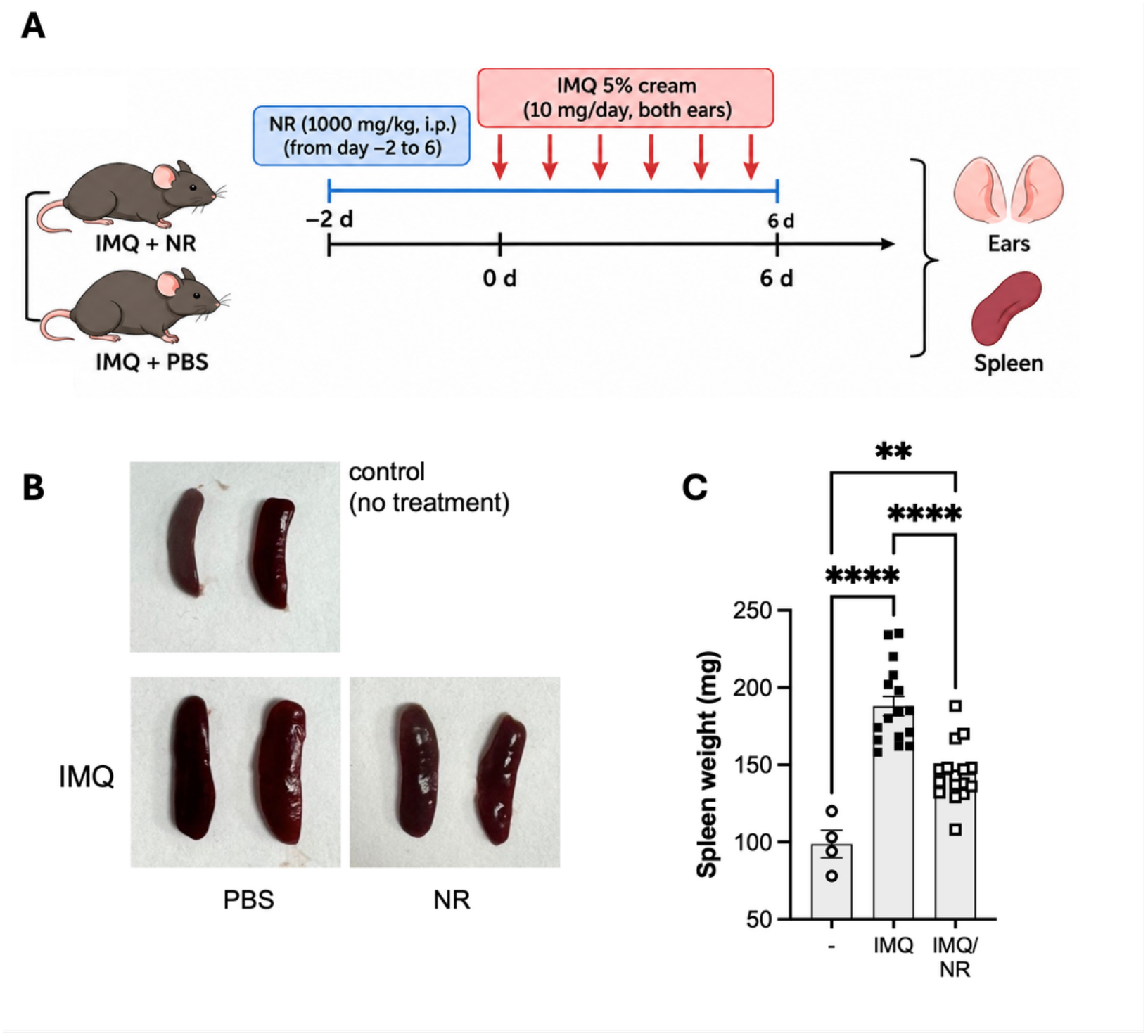
NR treatment attenuates IMQ-induced splenomegaly. (**A**) Experimental design: psoriasis-like skin inflammation was induced by daily topical application of imiquimod (IMQ). (**B**) Representative images of spleens from control, IMQ-treated, and IMQ+NR-treated mice. (**C**) Quantification of spleen weight. IMQ treatment induced a marked increase in spleen size, reflecting systemic inflammation, whereas NR supplementation significantly reduced splenomegaly, although spleen weight remained higher than in untreated controls. Data are shown as mean ± SEM, n=16. Statistical analysis was performed by one-way ANOVA followed by Tukey’s multiple comparisons test. *p < 0.05, **p < 0.01, ***p < 0.001, ****p < 0.0001.

Consistent with the systemic inflammatory response induced by IMQ, mice in the IMQ group exhibited a marked increase in spleen size compared with untreated controls.

Mean spleen size increased from 98.8 ± 18.8 mg in control animals to 187.9 ± 25.5 mg in IMQ-treated mice (p < 0.001). Administration of NR significantly reduced IMQ-induced splenomegaly (**Fig. 2B-C**), resulting in a mean spleen size of 144.3 ± 18.6 mg (p < 0.001 vs. IMQ). Spleen size in NR-treated mice remained intermediate between that of IMQ-treated and untreated control animals.

These findings indicate that NR treatment partially attenuated IMQ-induced splenomegaly, consistent with a reduction in the systemic inflammatory response.

### NR treatment reduces imiquimod-induced epidermal thickening and inflammation

Having observed that NR attenuated IMQ-induced splenomegaly, we next investigated whether NR treatment ameliorated the characteristic cutaneous manifestations of IMQ-induced psoriasiform inflammation, focusing on epidermal hyperplasia and inflammatory cell infiltration.

Epidermal thickness was measured to assess keratinocyte hyperplasia. IMQ treatment markedly increased epidermal thickness from 10.2 ± 1.8 μm in untreated mice to 41.5 ± 5.8 μm. NR treatment reduced epidermal thickness to 31.4 ± 2.4 μm (**Fig. 3A**). After accounting for basal epidermal thickness in untreated mice, this corresponds to an approximately 32% reduction in IMQ-induced epidermal thickening. The difference in epidermal thickness between IMQ-treated and NR-treated mice was statistically significant (unpaired two-tailed Student’s t-test, p < 0.001). These findings indicate that the treatment effectively attenuated IMQ-induced epidermal hyperplasia, a hallmark feature of psoriasiform skin inflammation.

**Figure 3.**
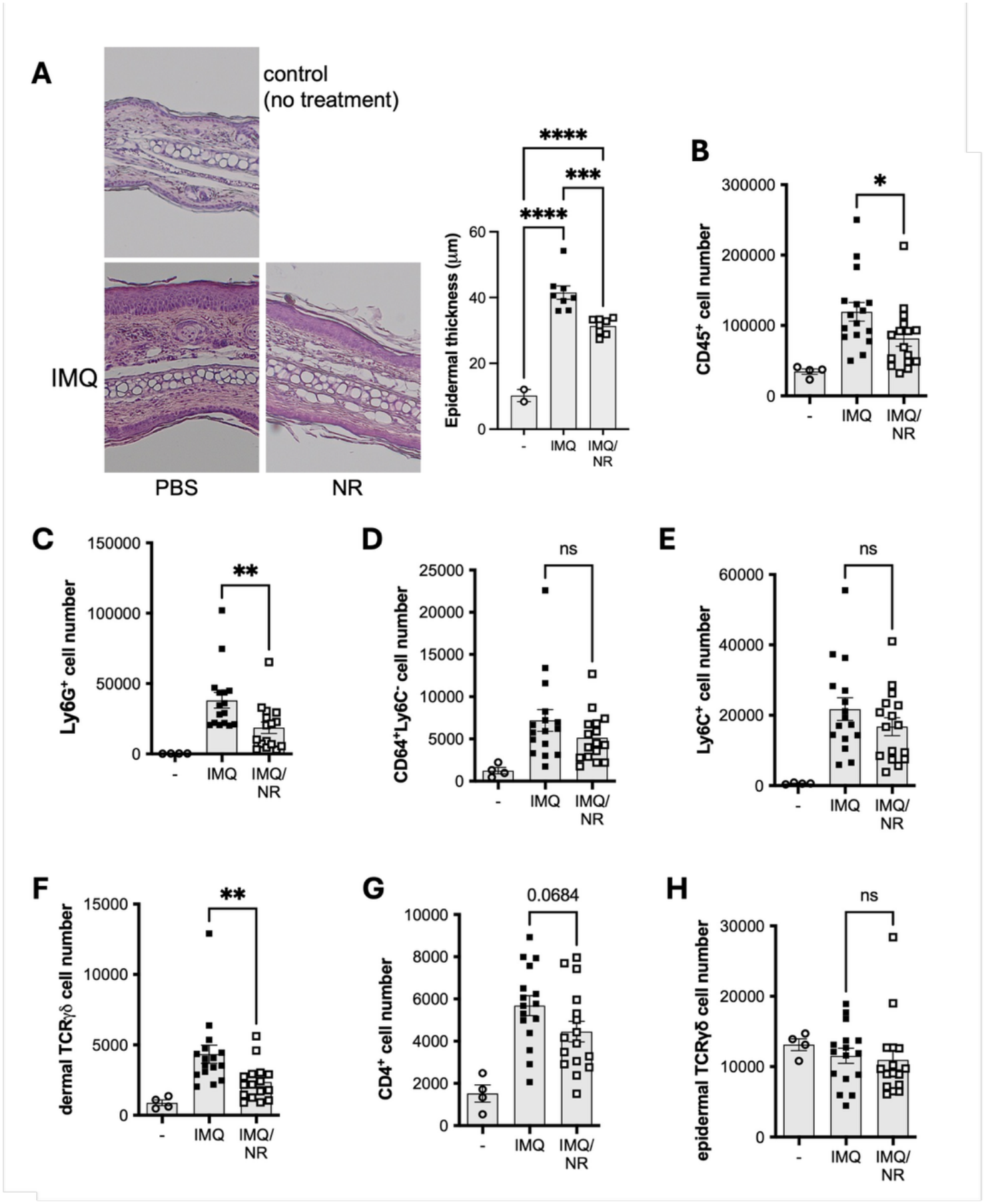
NR treatment reduces IMQ-induced epidermal hyperplasia and alters cutaneous immune cell populations. (**A**) Representative hematoxylin and eosin (H&E)-stained ear skin sections from control, IMQ-treated, and IMQ+NR-treated mice. Quantification of epidermal thickness. NR treatment significantly reduced IMQ-induced epidermal hyperplasia compared with IMQ-treated mice. Data are shown as mean ± SEM, n = 8. Statistical analysis was performed by one-way ANOVA followed by Tukey’s multiple comparisons test. (**B**–**G**) Flow cytometric analysis of inflammatory cell populations in ear skin. Representative gating strategy and quantification of CD45^+^cells (**B**), Ly6G^+^ neutrophils (**C**), CD64^+^Ly6C^+^ inflammatory monocyte/macrophages (**D**), CD64^+^Ly6C^-^ resident macrophages (**E**), dermal TCRγδ T cells (**F**), CD4^+^ T cells (**G**) and epidermal TCRγδ T cells (**H**). NR treatment reduced infiltration of both myeloid and lymphoid populations, with the most pronounced effect observed in neutrophils and dermal TCRγδ, which were restored to levels approaching those of untreated controls. Data are presented as mean ± SEM, n = 16. *p < 0.05, **p < 0.01, ***p < 0.001, ****p < 0.0001.

To further characterize the effects of NR on cutaneous inflammation, we quantified immune cell populations infiltrating the skin. NR treatment significantly reduced the total number of CD45^+^ leukocytes in the skin (**Fig. 3B**), indicating an overall reduction in immune cell infiltration. NR treatment also reduced both myeloid and lymphoid populations. Among myeloid cells, Ly6G^+^ neutrophils were decreased (**Fig. 3C**), while CD64^+^Ly6C^+^ inflammatory monocyte–macrophages and resident macrophages were largely unaffected (**Fig. 3D-E**). The reductions were more pronounced among lymphoid populations, particularly dermal TCRγδ T cells (**Fig. 3F**), whose numbers approached those observed in untreated animals, and CD4^+^ T cells (**Fig. 3G**). Notably, epidermal TCRγδ T cells showed little change following IMQ administration and were consequently not significantly affected by NR treatment (**Fig. 3H**).

Overall, these findings indicate that NR treatment ameliorates IMQ-induced skin inflammation by reducing epidermal hyperplasia and limiting the accumulation of inflammatory immune cells, as reflected by a reduction in total CD45^+^ leukocytes and in several specific myeloid and lymphoid populations.

## DISCUSSION

We and others previously showed that nicotinamide riboside (NR) exerts anti-inflammatory effects in myeloid and lymphoid cells through both sirtuin-dependent and sirtuin-independent mechanisms (Traba et al., 2015; Wu et al., 2022, 2024). By increasing NAD^+^ availability, NR has the potential to modulate immune-cell function and inflammatory responses through multiple, potentially complementary pathways. In line with this, our previous work demonstrated that NR reduced IL-17 production by activated CD4^+^ T cells isolated from healthy individuals and patients with psoriasis (Han et al., 2023), supporting a role for NAD^+^ metabolism in the regulation of Th17-associated inflammation. However, the effects of NR on psoriasis disease activity *in vivo* remained unexplored.

Here, we extend these findings by showing that NR attenuates IL-17A release by mouse CD4^+^ T helper 17 cells *ex vivo* and reduces IL-17A-associated inflammation *in vivo* in a mouse model of imiquimod-induced psoriasiform skin inflammation. Notably, NR did not significantly affect IFN-γ or IL-4 levels in mouse CD4^+^ T-cell cultures, suggesting that its effects are not associated with a generalized suppression of T-cell cytokine production but preferentially affect IL-17A-associated responses under the conditions examined. This selectivity is consistent with the prominent role of the IL-17 axis in imiquimod-induced skin inflammation, although the contributions from other inflammatory pathways cannot not be excluded.

Our previous studies have also provided insights into the mechanisms underlying the immunomodulatory effects of NR. In human CD4^+^ T cells, we found that NR suppressed Th17 responses in association with NRF2-dependent antioxidant responses, reduced ROS levels and changes in arginine and fumarate metabolism (Han et al., 2023). More recently, the involvement of SLIT2–ROBO1 signaling in the effects of NR on Th17 responses was described (Han et al., 2026), highlighting the contribution of dermal fibroblast–T-cell interactions. Together, these findings support the idea that NR can modulate T-cell inflammatory responses through multiple pathways, potentially contributing to its effects on cutaneous inflammation observed in the present study.

Dermal γδ17 cells are important contributors to IL-17-driven inflammation in murine models of psoriasis-like disease. Following imiquimod treatment, these cells produce substantial amounts of IL-17A and promote neutrophil recruitment and cutaneous inflammation (Ramírez-Valle et al., 2015; Borroto et al., 2022). Interestingly, although IL-17-producing γδ T cells have also been identified in human psoriatic lesions (Cai et al., 2011), αβ T cells appear to constitute the predominant T-cell population associated with human disease (Matos et al., 2017). Consistent with the attenuation of IL-17A-associated inflammation *ex vivo*, NR treatment reduced the abundance of dermal γδ17 cells, neutrophil infiltration, and the overall accumulation of CD45^+^ leukocytes in the skin. Together, these findings indicate that NR ameliorates imiquimod-induced psoriasiform inflammation and is associated with reduced accumulation of immune-cell populations implicated in IL-17-associated pathology. However, whether the reduction in dermal γδ17 cells reflects direct effects on their activation, proliferation, survival, or recruitment, or occurs secondarily to changes in the local inflammatory environment, remains to be established.

Importantly, the *in vivo* effects of NR are likely to extend beyond its cell-intrinsic effects identified in isolated CD4^+^ T-cell cultures and to involve additional mechanisms operating within the cutaneous inflammatory milieu. In our previous study, NR attenuated NLRP3 inflammasome activation and reduced IL-1β release in human PBMCs and monocytes (Traba et al., 2015), raising the possibility that NR also modulates innate immune signals that support IL-17-associated responses in the skin. Because IL-1β contributes to the activation and IL-17 production of dermal γδ17 cells, reduced IL-1β availability could represent one mechanism contributing to the decreased accumulation of these cells in NR-treated mice. Although this possibility remains to be directly tested in the present model, it provides a plausible link between the effects of NR on innate immune-cell function and the attenuation of IL-17-associated cutaneous inflammation. Thus, the *in vivo* activity of NR may reflect complementary effects on T-cell responses and the inflammatory signals that shape the local immune environment.

In conclusion, NR attenuates imiquimod-induced psoriasiform skin inflammation in mice, accompanied by reduced epidermal thickening and decreased accumulation of inflammatory leukocyte populations. These findings extend our previous observations on the immunomodulatory effects of NR and support further investigation of NAD^+^ metabolism as a potential target for modulating IL-17-associated skin inflammation. Whether these effects translate into clinical benefits in patients with psoriasis remains to be established.

## ACKNOWLEDGEMENTS

We thank the laboratory of Araceli del Arco, Beatriz Pardo and Jorgina Satrústegui (CBM) for their constant support and insightful discussion.

The laboratory of J.T. received funding from grants RYC2018-026050-I (funded by MICIU/AEI/10.13039/501100011033 and by ESF Investing in your future), PID2019-105665RA-I00 (funded by MICIU/AEI/10.13039/501100011033), and PID2022-136810OB-I00 (funded by MICIU/AEI/10.13039/501100011033 and ERDF/EU). This work also received support from grant RED2022-134927-T (MICIU/AEI; 10.13039/501100011033) and the NIH Office of Dietary Supplements (J.T.).

A.O. was supported by a contract funded by the Comunidad de Madrid (PEJ-2020-AI/BMD-17941). Y.C. was supported by predoctoral fellowship PRE2022-103127 (MICIU/AEI/10.13039/501100011033 and FSE+).

The Centro de Biología Molecular Severo Ochoa receives support through the Severo Ochoa Excellence grant (CEX2021-001154-S), funded by MICIU/AEI (10.13039/501100011033), and institutional support from the Ramón Areces Foundation.

The NR chemical powder was supplied by Chromadex Inc. Los Angeles, California, USA, under a Cooperative Research and Development Agreement (NIH – CTCR-19-004) to M.N.S. No funding for the research was supplied by Chromadex.

## Notes

### Competing Interest Statement

The authors have declared no competing interest.

